# Novel Linguistic Evaluation of Prefrontal Synthesis (LEPS) test measures prefrontal synthesis acquisition in neurotypical children and predicts high-functioning versus low-functioning class assignment in individuals with autism

**DOI:** 10.1101/467183

**Authors:** Andrey Vyshedskiy, Katarina Radi, Megan Catherine DuBois, Emma Mugford, Victoria Maslova, Julia Braverman, Irene Piryatinsky

**Author notes:** Corresponding author: Andrey Vyshedskiy, Ph.D., Boston University, Boston, USA.

## Abstract

In order to grasp the difference between “the cat on the mat” and “the mat on the cat,” understanding the words and the grammar is not enough. Rather it is essential to visually synthesize the cat and the mat together in front of the mind’s eye to appreciate their relations. This type of voluntary imagination, which involves juxtaposition of mental objects is conducted by the prefrontal cortex and is therefore called Prefrontal Synthesis (PFS). While PFS is essential for understanding of complex language, its acquisition has a strong experience-dependent critical period putting children with language delay in danger of never acquiring PFS and, consequently, not mastering complex language comprehension. In typical children, the timeline of PFS acquisition correlates with vocabulary expansion. Conversely, atypically developing children may learn hundreds of words but never acquire PFS. In these individuals, common tests of intelligence based on vocabulary assessment may miss the profound deficit in PFS. Accordingly, we developed a 5-minute test specific for PFS – Linguistic Evaluation of Prefrontal Synthesis or LEPS – and administered it to 50 neurotypical children, age 2 to 7 years (4.1±1.3) and to 23 individuals with impairments, age 8-21 years (16.4±3.0). All neurotypical children older than 4 years received the LEPS score 7/10 or greater indicating good PFS ability. Among individuals with impairments only 9 of 23 (39%) received the LEPS score 7/10 or greater. LEPS score was 90% correct in predicting high-functioning vs. low-functioning class assignment in individuals with autism, while full-scale IQ score was only 50% correct.

## Introduction

Language acquisition is a complex process that involves multiple cortical regions. Linking words with objects is the function of Wernicke’s area (Friederici, 2011), while interpreting the grammatical structure of a sentence and assigning word forms to a grammatical group (such as noun, verb, or preposition) is the function of Broca’s area (Friederici, 2011). Finally, combining objects from memory according to grammatically imposed rules into a novel mental image is the function of the lateral prefrontal cortex (LPFC) (Vyshedskiy, 2019; Vyshedskiy, Dunn, & Piryatinsky, 2017; Vyshedskiy, Mahapatra, & Dunn, 2017). The latter function is commonly called imagination. The term “imagination,” however, is ambiguous as it is regularly used to describe any unreal experience. For example, dreaming is often described as an imaginary experience. Dreaming though is not controlled by the LPFC (Braun et al., 1997; Siclari et al., 2017; Solms, 1997), since LPFC is inactive during sleep (Braun et al., 1997; Siclari et al., 2017) and patients whose LPFC is damaged do not notice change in their dreams (Solms, 1997). In order to distinguish the LPFC-dependent voluntary juxtaposition of mental objects from the LPFC-independent dreaming, we define the former as *Prefrontal Synthesis* or PFS (Vyshedskiy, 2019).

PFS is defined narrowly in order to distinguish it from other components of imagination, such as dreaming, mind-wondering, spontaneous insight, mental rotation, integration of modifiers, and also from other components of executive function, such as attention, impulse control, and working memory. PFS is not congruent to problem-solving, cognition, or fluid intelligence, as complex problems can often be solved via amodal completion (Gerbino & Salmaso, 1987; Weigelt, Singer, & Muckli, 2007), spontaneous insight (Salvi, Bricolo, Kounios, Bowden, & Beeman, 2016), integration of modifiers (Vyshedskiy, Dunn, et al., 2017) and other mechanisms, that do not involve LPFC-controlled juxtaposition of multiple objects (Vyshedskiy, 2019).

The notion about a special type of imagination different from dreaming and spontaneous insight, which is possibly unique to humans, has been entertained by many scientists. PFS has been described as “ability to invent fiction” (Harari, 2014), “episodic future thinking” (Atance & O’Neill, 2001), “mental scenario building” (Suddendorf & Redshaw, 2013), “mental storytelling” (Irwin, 2014), “internal mentation” (Andrews-Hanna, 2012), “mentally playing with ideas” (Diamond & Lee, 2011), “creative intelligence” (Fuster, 2003), “prospective memory” (Dobbs & Rule, 1987), “memory of the future” (Ingvar, 1985), “counterfactual thinking” (Roese, 1997). Traditionally, however, PFS ability was rolled into one of the more general abilities such as executive function, cognition, fluid intelligence, and working memory. None of those traits have a strong critical period since they can be improved well into adulthood (Jaeggi, Buschkuehl, Jonides, & Perrig, 2008). Only by defining PFS as a separate neurological mechanism, were we able to discover the strong critical period for PFS acquisition. Specifically, individuals who have not acquired PFS in early childhood cannot develop PFS later in life despite years of therapy (Vyshedskiy et al., 2019; Vyshedskiy, Mahapatra, et al., 2017). Strong critical periods are not unusual in central nervous system development. The most famous examples include monocular deprivation (Sherman & Spear, 1982), filial imprinting in birds (Bateson, 1979), and monaural occlusion (Knudsen, Knudsen, & Esterly, 1984). Note that PFS critical period is different from other language-related critical periods, such as phoneme tuning (Kral, 2013; Kuhl, Williams, Lacerda, Stevens, & Lindblom, 1992), grammar processing (Wartenburger et al., 2003), articulation control (Kim, Relkin, Lee, & Hirsch, 1997), and vocabulary acquisition (Snow & Hoefnagel-Höhle, 1978), which can be all significantly improved by training at any age (Kilgard & Merzenich, 1998; Tallal et al., 1996) and, therefore, have weak critical periods.

Less specific, more ambiguous definitions of PFS-like abilities water-down its strong critical period and undercut the analysis of language acquisition. E. g., theory-of-mind (ToM) is often included into PFS-like abilities. Similar to PFS, ToM acquisition has a critical period: deaf children who acquire formal sign language early, are significantly better at reasoning about mental states than language-delayed deaf children (Morgan & Kegl, 2006; Pyers & Senghas, 2009). ToM, however, improves at any age when individuals learn mental state vocabulary — particularly linguistic forms for verbs such as ‘think’ and ‘know’(Pyers & Senghas, 2009). Therefore, ToM has a weak critical period and shall not be merged with PFS that has a strong critical period.

Mental rotation and integration of modifiers are often also defined together with PFS since all three voluntary imagination processes are controlled by the LPFC. Similar to PFS, acquisition of mental rotation and integration of modifiers have critical periods (Martin, Senghas, & Pyers, 2013; Pyers, Shusterman, Senghas, Spelke, & Emmorey, 2010). However, both mental rotation and integration of modifiers, can be acquired in adulthood and therefore have weak critical periods (Curtiss, 1977; Grimshaw, Adelstein, Bryden, & MacKinnon, 1998). Accordingly, for the purposes of language acquisition, PFS must be considered separately from mental rotation and integration of modifiers.

Similar to other traits with strong critical periods – monocular deprivation, filial imprinting in birds, and monaural occlusion – PFS cannot be acquired in adulthood. Its neural infrastructure has to be laid down in early childhood. Perhaps this neural infrastructure is related to cortical functional specialization established through competition mechanisms similar to that of monocular deprivation (Ferjan Ramirez et al., 2013; Hinkley et al., 2016) and fine-tuning of long frontoposterior fibers, such as arcuate fasciculus and superior longitudinal fasciculus (Wilson et al., 2011), connecting these highly specialized cortical areas. The exact mechanism of the strong critical period for PFS acquisition remains to be determined.

All children not involved in external and internal recursive conversations are vulnerable to PFS paralysis. Among individuals with Autism Spectrum Disorder (ASD), the symptoms of PFS deficiency are commonly described as *stimulus overselectivity, tunnel vision*, or *lack of multi-cue responsivity* (Lovaas, Schreibman, Koegel, & Rehm, 1971) and 30-40% of individuals with ASD experience the associated lifelong impairment in the ability to understand spatial prepositions and recursion (Fombonne, 2003). These individuals, commonly referred to as having low-functioning ASD, typically exhibit full-scale IQ below 70 (Beglinger & Smith, 2001; Boucher, Mayes, & Bigham, 2008) and usually perform below the score of 85 in non-verbal IQ tests (Boucher et al., 2008). In fact, PFS and the associated ability to understand spatial prepositions and recursion, may be the most salient differentiator between high-functioning and low-functioning ASD.

The ASD medical community is aware of this strong critical period, and there is a wide consensus that intense early intervention should be administered to children as soon as they are diagnosed with ASD (Dawson et al., 2010). The goals of speech language pathologists (SLP) and Applied Behavioral Analysis (ABA) therapists happen to be built around the construct of PFS, and therefore it is highly targeted in these treatments. SLPs commonly refer to PFS developing techniques as “combining adjectives, location/orientation, color, and size with nouns,” “following directions with increasing complexity,” and “building the multiple features/clauses in the sentence” (American Speech-Language-Hearing Association, 2016). In ABA jargon, these techniques are known as “visual-visual and auditory-visual conditional discrimination” (Axe, 2008; Eikeseth & Smith, 2013; Lowenkron, 2006; Michael, Palmer, & Sundberg, 2011), “development of multi-cue responsivity” (Lovaas, Koegel, & Schreibman, 1979), and “reduction of stimulus overselectivity” (Ploog, 2010).

Despite the widespread recognition of the importance of early development of PFS abilities, there is a lack of psychometric tests that have the ability to measure a child’s progress in acquisition of PFS. Tests that rely exclusively on a child’s vocabulary, e.g., Peabody Picture Vocabulary Test (PPVT-4) (Dunn & Dunn, 2007) and Expressive Vocabulary Test (EVT-2) (Williams, 1997), are inadequate gauge of PFS: atypical children can learn hundreds of words, but fail to acquire PFS. Tests that require subjects to point to complex pictures, such as TONI-4 (Brown, Sherbenou, & Johnsen, 1982, 1997), Wechsler Intelligence Scale for Children (WISC-V) (Wechsler, 1949), Clinical Evaluation of Language Fundamentals (CELF-5) (Wiig, Secord, & Semel, 2013), Preschool Language Scales (PLS-5) (Zimmerman, Steiner, & Pond, 2011, p. 5), and Raven’s Progressive Matrices (J. Raven, 1998; J. C. Raven, 1936), can be too abstract for some individuals and set them up for failure (see (Lin & Chiang, 2014; Maljaars, Noens, Scholte, & van Berckelaer-Onnes, 2012), and also subject Peter described below, who failed at pointing to picture answers, but succeeded in comparable items with tangible objects). Finally, the Token Test (A. De Renzi & Vignolo, 1962; E. De Renzi & Faglioni, 1978) is unnecessarily difficult lexically, grammatically, and memory-wise and, therefore, lacks sensitivity to the PFS ability.

Various research groups have noticed this inability of both verbal and non-verbal IQ tests to adequately measure PFS in participants with impairments and developed an assortment of idiosyncratic tests to assess PFS. E. g., Grimshaw *et al*. (1998) studied a 19-year-old man referred to as E.M. E.M. who was born profoundly deaf and grew up in a rural area where he was not exposed to any formal sign language (Grimshaw et al., 1998). He and his family used homesign, a system of gestures that allowed them to communicate simple commands, but lacked much of recursion. This is quite typical of families with deaf children and hearing parents who are isolated from a sign language community (Mayberry, 2002; Mayberry & Eichen, 1991; Morford, 2003; Morford & Hänel-Faulhaber, 2011). Instead of learning a formal sign language, they spontaneously develop a homesign system. At the age of 15, E.M. was fitted with hearing aids that corrected his hearing loss and he began to learn verbal Spanish. When Grimshaw *et al*. tested E.M. at age 19, his performance on simple linguistic tests was “reasonably good”, but his performance on more complex tests that included spatial prepositions and recursion was “very poor.” Grimshaw *et al*. reported that “even at the 34-month assessment, he [E.M.] had not mastered one of these prepositions, nor were his errors limited to related pairs (under vs. over, in front of vs. behind). His general strategy when performing this subtest [following a direction to ‘put the green box in the blue box’] was to pick up the two appropriate objects and move them through a variety of spatial arrangements, watching the examiner for clues as to which was correct” (Grimshaw et al., 1998).

One of the most extensive evaluations of the function that we call PFS was conducted by Susan Curtiss in her analysis of language in Genie, a young girl who was linguistically isolated until the age of 12.7 (Fromkin, Krashen, Curtiss, Rigler, & Rigler, 1974). Curtiss’ battery of tests included over 20 verbal tasks intended to measure the extent to which Genie understood different aspects of language, including spatial prepositions, singular and plural sentences, negations with *un,* active vs. passive verb tense, superlatives, comparatives, and *wh*-questions. For example, in a test intended to measure Genie’s understanding of singular vs. plural nouns, Curtiss would present Genie with two pictures – one with one balloon and another with multiple balloons – and ask her to point to the picture of the balloon or balloons. Similarly, in a test intended to measure Genie’s understanding of superlatives, Curtiss would give Genie a picture of five buttons, each varying in size. Genie would be asked to point to the *smallest* or *largest* button, thereby indicating an understanding of superlative language. All of Curtiss’ other tests were structured this same way: Genie would be presented with objects or a picture of objects, given a question with a verbal instruction on which object/image to select, and asked to point or select accordingly. Similar to E.M., Genie’s performance on simple tests was reasonably good, but her performance on more complex tests was very poor. Genie never learned to understand spatial prepositions, recursion, and active vs. passive verb tense, i.e. functions that rely on PFS.

Alexander Luria worked extensively with adult patients whose PFS ability was compromised following a brain lesion. He reports that “these patients had no difficulty grasping the meaning of complex ideas such as ‘causation,’ ‘development,’ or ‘cooperation.’ They were also able to hold abstract conversations. But difficulties developed when they were presented with complex grammatical constructions which coded logical relations. … Such patients find it almost impossible to understand phrases and words which denote relative position and cannot carry out a simple instruction like ‘draw a triangle above a circle.’ This difficulty goes beyond parts of speech that code spatial relations. Phrases like ‘Sonya is lighter than Natasha’ also prove troublesome for these patients, as do temporal relations like ‘spring is before summer’. Additionally, patients with this type of lesion have no difficulty articulating words. They are also able to retain their ability to hear and understand most spoken language. Their ability to use numerical symbols and many different kinds of abstract concepts also remains undamaged. … Their particular kind of aphasia becomes apparent only when they have to operate with groups or arrangements of elements. If these patients are asked, ‘Point to the pencil with the key drawn on it’ or ‘Where is my sister’s friend?’ they do not understand what is being said. As one patient put it, ‘I know where there is a sister and a friend, but I don’t know who belongs to whom’” (Cole, Levitin, & Luria, 2014).

The multitude of idiosyncratic tests for PFS — stacking boxes used by Grimshaw *et al*. with subject E.M. (Grimshaw et al., 1998), spatial preposition tasks used by Curtiss with Genie (Fromkin et al., 1974), and mental reasoning (‘draw a triangle above a circle’) used by Luria with adults with LPFC lesions (Cole et al., 2014) — makes it difficult to compare results between different research groups. Accordingly, the purpose of this research was to develop a standardized test for assessment of PFS ability. We aimed to develop a test that measured PFS alone, exclusive of other components of imagination, such as spontaneous insight, mental rotation and integration of modifiers; a test that measured PFS directly without relying on vocabulary assessment. Most importantly, we wanted a test accessible to children with developmental and intellectual impairments including those who cannot comprehend tasks on paper. The resulting 5-minute PFS assessment was tested in a convenience sample of 50 typically developing children and in 23 individuals with impairments.

## Methods

The Linguistic Evaluation of Prefrontal Synthesis (LEPS) test is rooted in a set of common language comprehension items whereby the participants are required to follow verbal commands of increasing difficulty. The purpose of each item was to determine whether an individual could mentally combine several objects together, thereby indicating the level of overall PFS ability. All items were scored as either ***1:* participant has demonstrated an understanding of the item,** or ***0*: participant has not demonstrated an understanding of the item**.

The LEPS total score was calculated based on the number of items completed correctly. A total score of 10 indicated that a participant demonstrated an understanding of all items. Similarly, a participant who demonstrated an understanding of seven items would receive a total score of 7, a participant demonstrated an understanding of no items would receive a total score of 0, and so on.

The entire test was designed to take approximately 5 minutes to complete. A detailed description of each LEPS test item is provided below.

### 1. Integration of modifier

Integration of modifiers in a single object requires the participants to integrate a noun and an adjective. Participants were asked to select an object (e.g. *long red* straw) placed among several decoy objects including other *red* shapes (Lego pieces, small *red* animals) and *long/short* straws of other colors, thus forcing the participant to notice and integrate color, size and object. Colored straws were obtained from https://www.amazon.com/gp/product/B0721B4BJJ.

Prior to completing this item, participants were asked to point to and name the color of various objects to confirm that they understand the word for specific colors. Participants were then asked to complete four tasks in which colors, sizes, and nouns were varied randomly (Table 1, ‘Task examples’). Participants needed to answer correctly at least 3 out of 4 tasks (75% accuracy) to receive a score of *1* for this item. This 75% accuracy threshold was chosen to accommodate possible lapses in attention. With six colors, two sizes and three nouns, the probability of answering 75% of tasks correctly by chance is 0.004%. Thus, participants who made 1 error out of 4 tasks were highly unlikely to use the trial-and-error method and, therefore, demonstrated general understanding of the item.

**Table 1.**
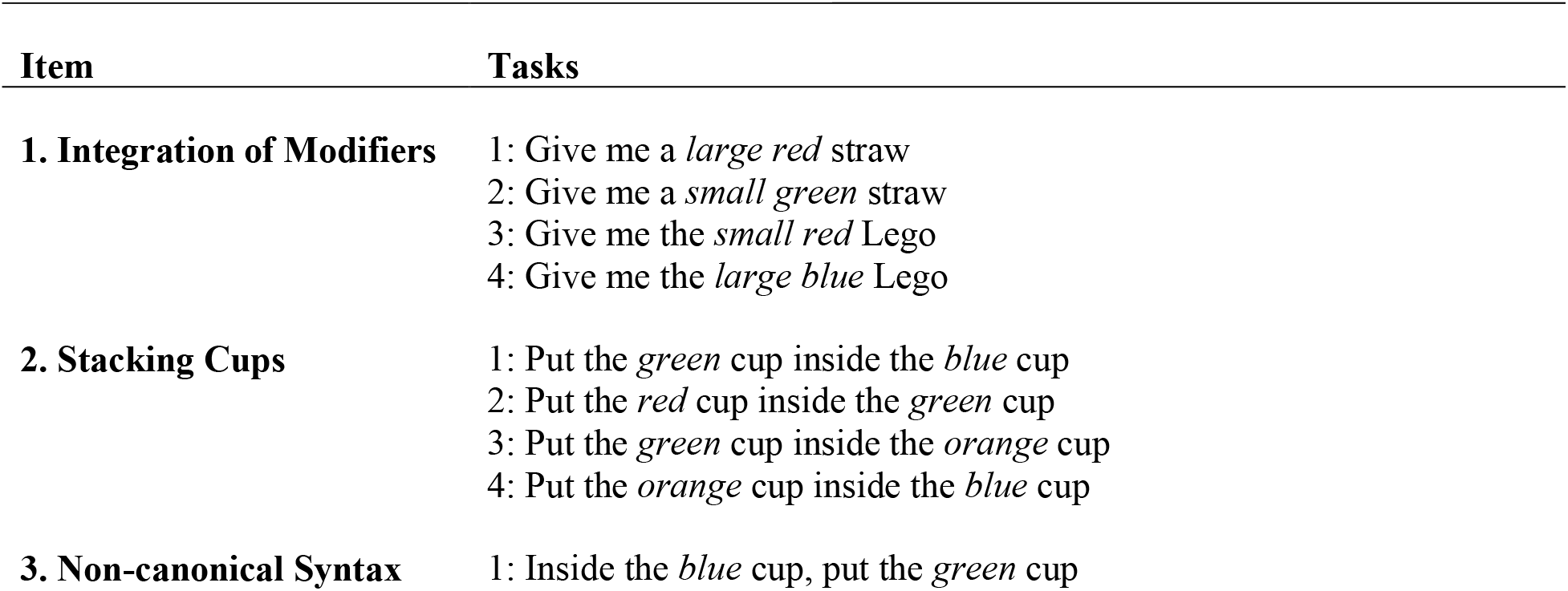
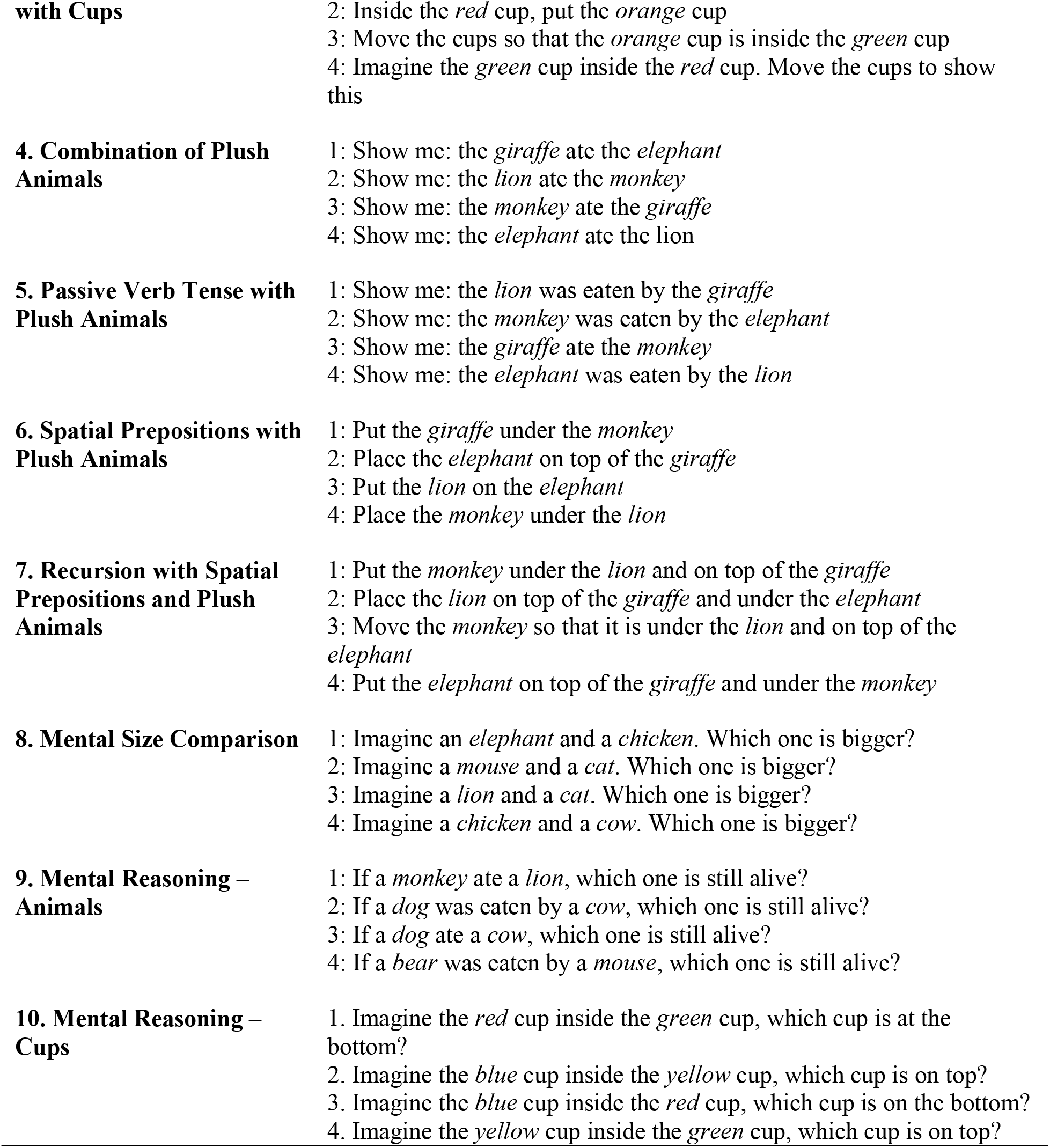
LEPS items and example questions.

### 2. Stacking cups

A set of colored cups (https://www.amazon.com/dp/B00GIPIM1U) was used for this test. The purpose of this task was to determine whether participants could properly arrange two cups, based on verbal instructions. Before the test, participants were given a demonstration of how to “put the *blue* cup inside the *red* cup” and, if necessary, were helped to stack the cups correctly. This training session with the *blue* and *red* cups was repeated while randomly switching the cup’s order until the participant was able to stack the correct cups on their own with no errors. Once subjects were comfortable stacking the two training cups, they were asked to stack four cups of various color combinations (Table 1, ‘Tasks’ column). Once the cups were stacked, each task was recorded as correct or incorrect. After each task, the tester encouraged the child by saying “Good job,” but no feedback was given concerning correctness of the answer in order to prevent the child from memorizing the answers.

Participants needed to answer correctly at least 3 out of 4 tasks (75% accuracy) to receive a score of *1* for this item. With four cup colors, the probability of answering 75% of tasks correctly by chance is 0.01%.

### 3. Non-canonical syntax with stacking cups

The directions for stacking cups were varied syntactically from the previous item. E. g., participants were instructed: “inside the blue cup, put the green cup” or “into the red cup, put the green cup.” This was intended to be a more difficult item than a canonical instruction like “put the green cup inside the blue cup.” Variation in syntax reduced the possibility that participants automatically remembered the instructions they were previously trained on in their language therapy and used this information to complete the item.

Participants needed to answer correctly at least 3 out of 4 tasks (75% accuracy) to receive a score of *1* for this item. With four cup colors, the probability of answering 75% of tasks correctly by chance is 0.01%.

### 4. Combination of plush animals

For this item, a set of puppet-like plush animals (giraffe, lion, elephant, and monkey https://www.amazon.com/gp/product/B075KRKPQ7) were laid on a flat surface. Each participant was asked to name the animals to confirm basic knowledge of animal names. Participants were shown an example of what it would look like if “the lion ate the monkey.” This was demonstrated by pushing the monkey puppet inside of the lion puppet. For this item, participants were instructed to manipulate puppets to show the experimenter what it would look like if “the elephant ate the lion” or if “the lion ate the elephant” and other similar variations.

Identically to all other items, at least 75% accuracy was required to earn a score of *1.* With four animals, the probability of answering 75% of tasks correctly by chance is 0.01%.

### 5. Passive verb tense with plush animals

This item used the same plush animals, but sought to measure whether the participant could still correctly position one animal inside of another when the directions were given in passive verb tense. For example, participants were prompted with the directions: “the giraffe was eaten by the lion.” This decreased the likelihood that participants could follow a rigid routinized algorithm (the first animal is the predator; the second animal is the prey). Since the positions of the predator and the prey in a sentence vary randomly, participants are more likely to actually imagine which animal ate the other.

Again, at least 75% accuracy was required to earn a score of *1.* With four animals, the probability of answering 75% of tasks correctly by chance is 0.01%.

### 6. Spatial prepositions with Plush Animals

In this item, participants were instructed to maneuver the plush animals according to the spatial prepositions *on top of* and *under*. Before the test, participants were given a demonstration of how to “put the monkey *on top of* and *under* the lion” and, if necessary, were helped to stack the animals correctly. This training session with the monkey and lion was repeated while randomly switching the order of animals until the participant was able to stack the animals on their own with no errors. Once subjects were comfortable stacking the two training animals, participants were asked to show “the giraffe under the monkey,” or “the elephant on top of the giraffe.” The pair containing monkey and lion was not used in the actual test.

The spatial prepositions *behind* and *in front of* were not used to avoid confusion of whether the perspective was from the experimenter or the participant.

Participants needed to answer correctly at least 3 out of 4 tasks (75% accuracy) to receive a score of 1 for this item. With four animals, the probability of answering 75% of tasks correctly by chance is 0.01%.

### 7. Recursion with spatial prepositions

Recursion with spatial prepositions was also used to verbally indicate the position of the plush animals. Participants were instructed in the following way: “show me: the monkey is under the lion and on top of the giraffe.” The instructions always used the middle animal as the point of reference so that the participant had to mentally integrate both aspects of the direction to arrange the animals.

Identically to all other items, at least 75% accuracy was required to earn a score of *1.* With four animals, the probability of answering 75% of tasks correctly by chance is 0.00007%.

### 8. Mental size comparison

This item included verbal questions in which participants were asked to tell the tester which animal was bigger than the other. For example, the participants were asked “which animal is bigger: the elephant or the chicken?” or “the cat or the mouse?” or “the cat or the lion?” In this item, participants had to determine which animal was bigger than the other by using their own mental representations of animals, without the use of physical representations. Again, at least 75% accuracy was required to earn a score of *1.* The probability of answering 75% of tasks correctly by chance is 25%.

### 9. Mental reasoning - animals

In the final two items, participants were asked to synthesize multiple pieces of information to solve simple mental reasoning tasks. No tangible objects were used as representation. In this item, the task was about animal predation. For example, the prompt could be: “if the monkey ate a snake, who is alive?” or “if a lion was eaten by a snake, who is alive”? Instructions for this item included both passive and active verb tenses, which added an extra level of difficulty. Identically to all other items, at least 75% accuracy was required to earn a score of *1.* The probability of answering 75% of tasks correctly by chance is 25%.

### 10. Mental reasoning - cups

In the final item, the task required participants to imagine stacking cups. For example, the prompt could be: “imagine the red cup inside the green cup, which cup is at the bottom?” or “imagine the blue cup inside the yellow cup, which cup is on top”? Again, at least 75% accuracy was required to earn a score of *1.* The probability of answering 75% of tasks correctly by chance is 25%.

### Neurotypical participants

A convenience sample of neurotypical participants were obtained for this study by approaching parents of young children in local parks and asking if they would be willing to let a researcher administer the test to their child. The majority of neurotypical participants were obtained from parks in an affluent suburb of Boston, indicating that the convenience sample represents a relatively privileged population. The data presented in this manuscript include everyone, who agreed to be tested, except one child whose parents indicated that he has suspected ASD and two children whose parents indicated that they had a significant developmental delay. All participants’ caregivers consented to anonymized data analysis and publication of the results, and were present during test administration. The mean age of neurotypical participants was 4.1±1.3 (range 2-7) and 44% of them were male.

### Participants with impairments

LEPS test was also administered to 23 individuals with ASD or other developmental and intellectual impairments. To select the participants, we reached out to all parents of students in four classes at the Bancroft school in Mt. Laurel, NJ (27 students): two classes from high-school program (age range: 14-18) and two classes from post-high school transitional program (age range: 18-21). All students whose parents have signed parental consent (20 participants, 74%) were administered LEPS test and are included in the study. In addition, we describe three convenience participants used in LEPS development. The mean age of participants with impairments was 16.4±3.0 years, range: 8-21 years, Table 2. The mean full-scale IQ=56±12, verbal IQ=57±11, nonverbal IQ=65±14. 91% of participants with impairments were male. Five participants in this study were not able to communicate using vocal speech. They responded via Alternative Augmentative Communication devices (4 participants) and American Sign Language (1 participant).

**Table 2.** Participants with impairments.

| **ID** | **Age**<br>**(years)** | **Sex** | **Full-**<br>**scale**<br>**IQ** | **Verbal**<br>**IQ** | **Non-**<br>**verbal**<br>**IQ** | **Communication**<br>**abilities** | **Diagnosis** | **Class**<br>**assignment** | **LEPS**<br>**total** |
| --- | --- | --- | --- | --- | --- | --- | --- | --- | --- |
| **1** | 18.2 | m | 40 |  | 47 | nonverbal | ASD,<br>ADHD | Low-func. | **0** |
| **2** | 17.1 | m |  |  |  | **verbal** | ASD |  | **2** |
| **3** | 15.1 | m | low |  |  | **verbal** | ASD | Low-func. | **2** |
| **4** | 16.3 | f | low | 56 |  | nonverbal | ASD | Low-func. | **2** |
| **5** | 15.9 | m | 62 |  |  | **verbal** | ASD | Low-func. | **3** |
| **6** | 14.6 | m |  |  |  | nonverbal | ASD | Low-func. | **3** |
| **7** | 14.7 | m |  |  |  | **verbal** | ASD | **High-func.** | **3** |
| **8** | 15.1 | m |  |  |  | **verbal** | ASD | Low-func. | **4** |
| **9** | 21.3 | f | 45 | 47 | 65 | **verbal** | ASD, OCD | Low-func. | **4** |
| **10** | 18.2 | f | 52 | 63 | 58 | **verbal** | ASD,<br>ADHD,<br>Epilepsy | Low-func. | **4** |
| **11** | 15.4 | m |  |  |  | nonverbal | ASD | Low-func. | **5** |
| **12** | 18.7 | f | 43 | 43 | 44 | **verbal** | ASD,<br>cerebral<br>palsy | Low-func. | **5** |
| **13** | 18.6 | m | 51 | 66 | 56 | **verbal** | ADD,<br>emotionally<br>disturbed | Low-func. | **6** |
| **14** | 16.0 | m |  |  | 72 | **verbal** | ASD | **High-func.** | **6** |
| **15** | 19.6 | m |  |  |  | **verbal** | ASD | **High-func.** | **7** |
| **16** | 20.8 | m | 52 | 68 | 60 | **verbal** | ASD,<br>ADHD | **High-func.** | **8** |
| **17** | 17.4 | m | 78 |  |  | **verbal** | ASD,<br>ADHD | **High-func.** | **9** |
| **18** | 17.0 | m | 59 |  |  | **verbal** | ASD | **High-func.** | **9** |
| **19** | 15.8 | m |  |  |  | **verbal** | ASD | **High-func.** | **9** |
| **20** | 7.6 | m | 79 |  | 74 | **verbal** | ADHD |  | **9** |
| **21** | 17.3 | m | 52 |  |  | **verbal** | ASD | **High-func.** | **10** |
| **22** | 15.9 | m | 63 | 70 | 74 | **verbal** | ASD | **High-func.** | **10** |
| **23** | 10.3 | m |  |  | 74 | nonverbal | ASD |  | **10** |

## Results

Fifty neurotypical and 23 participants with impairments were tested using the LEPS test. Each item aimed to determine whether an individual could integrate and imagine several objects together, thereby indicating the level of overall PFS abilities. Using variations in sentence structure for each verbal instruction, every subject was asked to complete four tasks in each item. All items were scored as either *1:* participant has demonstrated an understanding, or *0*: participant has not demonstrated an understanding.

### 1. Integration of modifier

The purpose of this item is to determine whether an individual is able to integrate two different properties of an object (e.g., *give me the small red straw –* find both the small straw and the red straw, thus integrating two properties). All neurotypical children over the age of four and 91% of participants with impairments demonstrated an understanding of this item, indicating that each understood the basic properties of an object and could combine nouns and adjectives to select the correct object (Tables 3 and 4).

**Table 3.** Neurotypical children % correct in each LEPS item in each age group.

| **Age**<br>**(years)** | **Item**<br>**1** | **Item**<br>**2** | **Item**<br>**3** | **Item**<br>**4** | **Item**<br>**5** | **Item**<br>**6** | **Item**<br>**7** | **Item**<br>**8** | **Item**<br>**9** | **Item**<br>**10** | **LEPS**<br>**total** |
| --- | --- | --- | --- | --- | --- | --- | --- | --- | --- | --- | --- |
| **2-3** | 27% | 33% | 7% | 7% | 0% | 0% | 0% | 20% | 0% | 0% | **0.9±1.4** |
| **3-4** | 100% | 83% | 50% | 67% | 50% | 33% | 0% | 83% | 33% | 25% | **5.3±2.2** |
| **4-5** | 100% | 100% | 88% | 100% | 75% | 88% | 63% | 100% | 100% | 75% | **8.9±1.1** |
| **5-6** | 100% | 100% | 100% | 100% | 100% | 100% | 60% | 100% | 80% | 80% | **9.2±0.8** |
| **6-7** | 100% | 100% | 100% | 100% | 100% | 100% | 40% | 100% | 100% | 100% | **9.4±0.5** |

**Table 4.** Participants with impairments performance in each LEPS item.

| **ID** | **Item 1** | **Item 2** | **Item 3** | **Item 4** | **Item 5** | **Item 6** | **Item 7** | **Item 8** | **Item 9** | **Item 10** | **LEPS total** |
| --- | --- | --- | --- | --- | --- | --- | --- | --- | --- | --- | --- |
| **1** | 0 | 0 | 0 | 0 | 0 | 0 | 0 | 0 | 0 | 0 | **0** |
| **2** | 1 | 1 | 0 | 0 | 0 | 0 | 0 | 0 | 0 | 0 | **2** |
| **3** | 1 | 0 | 0 | 0 | 0 | 0 | 0 | 1 | 0 | 0 | **2** |
| **4** | 1 | 0 | 0 | 0 | 0 | 0 | 0 | 1 | 0 | 0 | **2** |
| **5** | 1 | 1 | 0 | 0 | 0 | 0 | 0 | 1 | 0 | 0 | **3** |
| **6** | 1 | 0 | 0 | 0 | 0 | 0 | 0 | 0 | 1 | 1 | **3** |
| **7** | 0 | 1 | 1 | 0 | 1 | 0 | 0 | 0 | 0 | 0 | **3** |
| **8** | 1 | 1 | 0 | 1 | 0 | 0 | 0 | 0 | 0 | 1 | **4** |
| **9** | 1 | 1 | 0 | 1 | 0 | 0 | 0 | 1 | 0 | 0 | **4** |
| **10** | 1 | 1 | 0 | 0 | 0 | 1 | 0 | 1 | 0 | 0 | **4** |
| **11** | 1 | 0 | 1 | 1 | 0 | 0 | 0 | 1 | 1 | 0 | **5** |
| **12** | 1 | 1 | 0 | 0 | 1 | 1 | 0 | 1 | 0 | 0 | **5** |
| **13** | 1 | 1 | 0 | 1 | 0 | 1 | 0 | 1 | 0 | 1 | **6** |
| **14** | 1 | 1 | 0 | 0 | 1 | 1 | 0 | 1 | 1 | 0 | **6** |
| **15** | 1 | 1 | 1 | 1 | 1 | 1 | 0 | 1 | 0 | 0 | **7** |
| **16** | 1 | 1 | 1 | 1 | 1 | 1 | 0 | 1 | 0 | 1 | **8** |
| **17** | 1 | 1 | 1 | 1 | 1 | 1 | 0 | 1 | 1 | 1 | **9** |
| **18** | 1 | 1 | 0 | 1 | 1 | 1 | 1 | 1 | 1 | 1 | **9** |
| **19** | 1 | 1 | 1 | 1 | 1 | 1 | 1 | 1 | 0 | 1 | **9** |
| **20** | 1 | 1 | 1 | 1 | 0 | 1 | 1 | 1 | 1 | 1 | **9** |
| **21** | 1 | 1 | 1 | 1 | 1 | 1 | 1 | 1 | 1 | 1 | **10** |
| **22** | 1 | 1 | 1 | 1 | 1 | 1 | 1 | 1 | 1 | 1 | **10** |
| **23** | 1 | 1 | 1 | 1 | 1 | 1 | 1 | 1 | 1 | 1 | **10** |
| **% correct** | **91%** | **78%** | **43%** | **57%** | **48%** | **57%** | **26%** | **78%** | **39%** | **48%** |  |

### 2. Stacking cups test

In this item, participants were tested on their ability to correctly stack two cups in the order instructed. For example, participants were instructed to “put the blue cup inside the green cup.” All neurotypical children over the age of four and 78% of participants with impairments demonstrated an understanding of this item.

### 3. Non-canonical syntax with stacking cups

The non-canonical syntax with stacking cups item was intended to measure whether participants could correctly order two cups when syntax deviated from a canonical structure: e.g. “inside the blue cup, put the red cup.” 96% of neurotypical children over the age of four and only 43% of participants with impairments demonstrated an understanding of this item.

### 4. Combination of plush animals

The purpose of this item is to determine whether participants could correctly determine which plush animal ate the other and arrange the animals accordingly based on verbal instructions, such as “the lion ate the monkey.” All neurotypical children over the age of four and only 57% of participants with impairments demonstrated an understanding of this item.

### 5. Passive verb tense with plush animals

The purpose of this item is to determine whether participants could still order the plush animals when the verb tense changed from active to passive (e.g. “the lion was eaten by the monkey.”) 91% of neurotypical children over the age of four and 48% participants with impairments demonstrated an understanding of this item.

### 6. Spatial prepositions with plush animals

The purpose of this item is to understand subjects’ ability to follow directions with spatial prepositions, e.g. “show me the monkey under the giraffe.” 96% of neurotypical children over the age of four and 57% participants with impairments demonstrated an understanding of this item.

### 7. Recursion with spatial prepositions

The purpose of this item is to determine participants’ ability to follow prepositional directions to determine where the animals should be located relative to each another, while the directions were always centered on the middle animal. For example, “the monkey is under the lion and on top of the giraffe.” 57% of neurotypical children over the age of four and 26% participants with impairments demonstrated an understanding of this item. This was clearly the most difficult item of the test.

### 8. Mental size comparison

The purpose of this item is to ask the participant to mentally visualize different animals in their mind and determine which one was bigger, e.g. “which animal is bigger, a lion or a cat?” All neurotypical children over the age of four and 78% participants with impairments demonstrated an understanding of this item.

### 9. Mental reasoning – animals

The purpose of this item is to assess whether participants are able to mentally imagine a scene in which one animal ate another, without physical representations of the animals. The instructions used both active and passive verb tenses: e.g., “if the snake ate the lion, who is alive” or “if the snake was eaten by the lion, who is alive.” 91% of neurotypical children over the age of four and 39% participants with impairments demonstrated an understanding of this item.

### 10. Mental reasoning – cups

The purpose of this item is to assess whether participants are able to mentally imagine the order of stacked cups without physical representations of them. 83% of neurotypical children over the age of four and 48% participants with impairments demonstrated an understanding of this item.

### Age of Prefrontal Synthesis acquisition in neurotypical children

Figure 1 summarizes the LEPS total score as a function of age in neurotypical children. Markers indicate individual children LEPS scores. Notice the exponential increase of the LEPS total score between the ages of 3 and 4.

**Figure 1.**
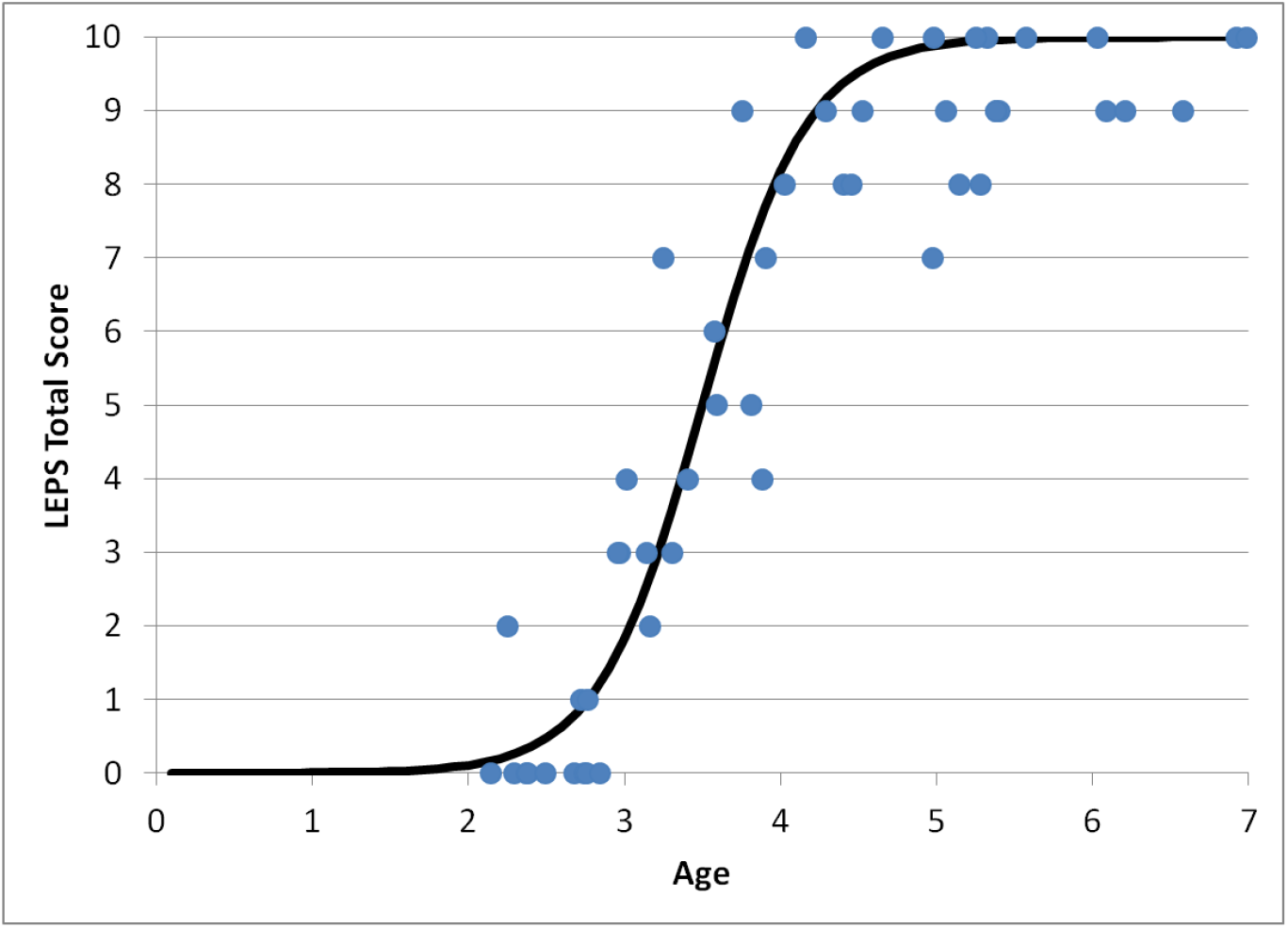
Scatter plot of the LEPS total score as a function of age in neurotypical children. Markers indicate LEPS total scores of individual children.

### Psychometric Characteristics of LEPS

#### Reliability

Reliability was assessed in neurotypical participants. Internal consistency was excellent (Cronbach’s alpha equals to 0.95), suggesting high reliability. All items demonstrated high (>0.5) item-total correlations. The LEPS test-retest reliability was evaluated by calculating a Pearson Correlation between the first administration of the LEPS and the re-administration of the LEPS to the same participants approximately 2 months (23 - 91 days) later (23 participants). The 2-month test-retest correlation coefficient for LEPS was r =0.96 (p < 0.001), revealing excellent LEPS long-term stability.

#### Validity

To demonstrate known group validity, the LEPS total score was compared to students’ class assignment. Like at many other educational institutions, at Bancroft, students are grouped into higher-level and lower-level classes. For the purposes of this assignment, Bancroft does not use any standardized metrics. IQ scores are not considered, “because teachers don’t have good experience with IQ scores - they seem to be inaccurate.” Rather, teachers and clinical teams assign students to classes based on their own experience and informal assessments.

Bancroft students who participated in this study came from four classes: 1) high-school program (age range: 14-18) higher-functioning class, 2) high-school program lower-functioning class, 3) transitional program (age range: 18-21) higher-functioning class, 4) transitional program lower-functioning class. Altogether, 11 participants came from lower-functioning classes and 9 participants came from higher-functioning classes, Table 2. LEPS score had excellent prediction power for students’ class assignment. All students with LEPS total score ≥7 were assigned by teachers to higher-functioning classes; all but two students with LEPS total score <7 were assigned by teachers to lower-functioning classes, resulting in LEPS class assignment prediction rate of 90% (18 correct predictions out of 20 students). The t-test demonstrated significant difference between mean LEPS scores for students in higher- vs. lower-functioning classes (3.3±1.7 vs. 7.9±2.3; t(18) = 5.11; p<0.0001). This supports high known-group validity of LEPS.

An additional evidence for known-group validity of LEPS comes from the comparison between neurotypical and atypical individuals. All neurotypical children older than 4 years received the LEPS score 7/10 or greater indicating good PFS ability. At the same time, among individuals with impairments only 9 of 23 (39%) received the LEPS score 7/10 or greater. The t-test demonstrated a significant difference between the mean LEPS score in neurotypical children older than 4 years old and participants with impairments (9.1±0.9 vs. 5.7±3.1, t(26) = 2.40; p<0.03).

At the same time LEPS demonstrates high discriminant validity when compared with the IQ test. LEPS was only moderately correlated with IQ scores. Specifically, the Pearson coefficients were as following: full-scale IQ=0.54, verbal IQ=0.34, nonverbal IQ=0.55. This shows that LEPS is conceptually different from IQ.

#### Inter-observer agreement

Inter-observer agreement was assessed in nine participants with impairments. Two observers independently scored every question in each of 10 items for the total of 40 questions per a test. The inter-observer agreement was very high as indicated by Kappa analysis with Kappa = 0.98 and the percent overall agreement = 99.2%.

## Discussion

Association of Wernicke’s and Broca’s areas with language is well-known. Less common is the realization that understanding of full language depends on the lateral prefrontal cortex (LPFC). Wernicke’s area primarily links words with objects (Friederici, 2011), Broca’s area interprets the grammar and assigns words in a sentence to a grammatical group such as noun, verb, or preposition (Friederici, 2011), but only the LPFC can synthesize the objects from memory into a novel mental image according to grammatically imposed rules (Vyshedskiy, Dunn, et al., 2017; Vyshedskiy, Mahapatra, et al., 2017). This latter function may be called *imagination*, but we prefer a more specific term, *Prefrontal Synthesis* (PFS) in order to distinguish this function from other components of imagination, such as dreaming, simple memory recall, spontaneous insight, mental rotation, and integration of modifiers (Vyshedskiy, 2019). PFS is defined as voluntary juxtaposition of mental objects.

PFS is essential for understanding sentences describing combinations of objects. E.g., the sentences “The dog bit my friend” and “My friend bit the dog” use identical words and grammar. Appreciating the misfortune of the first sentence and the humor of the second sentence depends on the LPFC ability to faithfully synthesize the two objects – the friend and the dog – into a novel mental image. Similarly, understanding of spatial prepositions such as *in, on, under, over, beside, in front of, behind* requires a subject to synthesize several objects in front of the mind’s eye. For example, the request “to put a green box {inside/behind/on top of} the blue box” requires an initial mental simulation of the scene, only after which is it possible to correctly arrange the physical objects. An inability to produce a novel mental image of the green box {inside/behind/on top of} the blue box would lead to the use of trial-and-error, which in majority of cases will result in an incorrect arrangement.

PFS completely depends on the intact LPFC and patients with damage to the LPFC often lose their PFS function (Baker et al., 1996; Christoff & Gabrieli, 2000; Duncan, Burgess, & Emslie, 1995; Fuster, 2008; A. Luria, 2012; Waltz et al., 1999). Fuster calls their condition “prefrontal aphasia” (Fuster, 2008) and Luria defines it as “frontal dynamic aphasia” (A. R. Luria, 1970). Fuster explains that “although the pronunciation of words and sentences remains intact, language is impoverished and shows an apparent diminution of the capacity to ‘prepositionize.’ The length and complexity of sentences are reduced. There is a dearth of dependent clauses and, more generally, an underutilization of what Chomsky characterizes as the potential for recursiveness of language.” (We prefer to refer to this condition as ‘PFS paralysis’ since aphasia is translated from Greek as “speechless” and these patients may not experience any speech deficit.)

Typically developing children acquire PFS naturally and their progress is obviated by their conversations. In children with language delay, monitoring PFS acquisition is much more tricky since in some individuals it can fall behind the simpler function of vocabulary acquisition creating a false sense of normalcy (Boucher et al., 2008; Hudry et al., 2010; Lovaas et al., 1971; Maljaars et al., 2012). It is not uncommon to observe the following developmental steps in individuals who acquire language with a significant delay: they start to understand some individual words and phrases, then develop understanding of recursion, and only after that they begin to verbally express themselves, first with individual words and then with complete sentences. The existing evaluations adequately assess the former (receptive vocabulary acquisition, Wernicke’s area) as well as the latter (expressive language development, Broca’s area), but, critically, miss to assess the middle step which heralds the LPFC function of PFS. Therefore, there is a substantial gap in the ability of the existing evaluation tools to faithfully measure an important part of child’s developmental progress.

The purpose of this research was to develop a test for PFS that is suitable for children, particularly those with language delay. PFS assessment in adults has a range of options, such as the Tower of London test (Shallice, 1982) and the mental 2-digit number multiplication (Zago et al., 2001). However, these tests are not applicable to young children, as they rely heavily on attention and working memory, which tax the PFC beyond abilities of most children, and knowledge of multiplication that is beyond the limits of young children who do not yet know arithmetic. Accordingly, we developed a 10-item Linguistic Evaluation of Prefrontal Synthesis (LEPS) scale and used it to assess PFS in 50 neurotypical children age 2 to 7 years (4.1±1.3) and in 23 individuals with impairments, age 8-21 years (16.4±3.0). The LEPS test exhibited excellent internal consistency, excellent inter-observer agreement, and excellent test-retest reliability. In neurotypical children the LEPS score increased exponentially from 3 to 4 years, Figure 1. Not a single child aged 3.1 years and younger received the LEPS score ≥7. All neurotypical children aged 4 years and older received the LEPS score ≥7 indicating PFS ability. Among individuals with impairments 9 of 23 (39%) received the LEPS score ≥7 indicating PFS ability and 14 of 23 (61%) received the LEPS score <7 indicating PFS paralysis or partial paralysis. In individuals with impairments LEPS score had excellent predictive power for student higher-functioning vs. lower-functioning class placement and weak correlation with IQ. In the following discussion, we describe our logic for LEPS format and individual items, as well as several notable observations from the development of the measure that provide insight into how and why LEPS can be used to test PFS in atypically developing children.

### Integration of modifiers is simpler than PFS

Integration of modifiers is the first item of the LEPS test. Neurologically, both functions, integration of modifiers and PFS, are controlled by the LPFC (Vyshedskiy, 2019). However, integration of modifiers only involves modification of neurons encoding a single object and, consequently, is simpler than the process of PFS which, by definition, involves combination of *several* objects (Vyshedskiy, Dunn, et al., 2017). In other words, integration of modifiers is not PFS, but a developmental precursor to PFS. All neurotypical participants over the age of three and 91% of participants with impairments demonstrated an understanding of this item. Compare that observation to the number of participants who demonstrated presence of PFS (defined as the LEPS total score of 7 or more): 77% of neurotypical participants over the age of three and only 39% of participants with impairments.

While item 1 does not evaluate PFS, it was included in LEPS for several reasons. First, it is useful for quick assessment of participant’s understanding of colors and sizes – essential elements used throughout the LEPS test. Second, in participants with PFS paralysis, it is useful to know if at least the precursor to PFS has been acquired. Third, the easy task of integration of modifiers is a convenient way to focus the participant on more difficult items. Finally, the integration of modifiers item can be used repeatedly throughout LEPS test to gauge participant’s attention.

### The LEPS test attempts to avoid routinized responses

The purpose of the remaining items 2 to 10 was to create a series of mental puzzles that varied syntactically from canonical instructions that could be routinized through long-term training. E. g., consider item 2 that instructed participants to “put the green cup inside the blue cup.” There are two ways to successfully complete this stacking cup instruction. One way to find the solution is to mentally synthesize a novel image of the green cup inside the blue cup, and then, after completing the mental simulation, arrange the physical objects to match the image in the mind’s eye. An alternative solution could be obtained algorithmically by following these steps: (1) lift the cup mentioned first; (2) insert it into the cup mentioned second. This type of algorithmic solution does not require PFS. It is a sort of automatic routinized action encoded in basal ganglia, akin to riding a bicycle, tying shoelaces, skiing, skating, stopping at a red light, or writing a signature.

All but one neurotypical children 3.8 years or older (96%) were able to demonstrate understanding of non-canonical instructions in item 3. On the contrary, out of 19 participants with impairments who were able to complete the canonical stacking cups task in item 2, only 10 (53%) were able to complete the same task under the condition of non-canonical word order in item 3. Failing participants with impairments usually selected the correct cups, but assembled them randomly (see Movie 1 for a video recording of a typical student). Their performance is consistent with highly routinized response. Most participants with impairments have received over 15 years of intensive language therapy and it is likely that their stacking cups routine has been automated through frequent ‘stacking cups’ training using canonical syntax only. Notably, most failing individuals completed each stacking movement fast, with no hesitation. Neurotypical children, on the other hand, normally paused to think while completing the same task, presumably to simulate the answer mentally.

Naturally, in a test for PFS, we wanted to avoid giving participants an opportunity to answer items algorithmically as much as possible. Theoretically, if we knew which tasks individuals were trained on, we could have avoided those tasks in the test items. However, it is not feasible for a formal test to avoid all tasks a participant could have been trained on. An alternative to this predicament would be to increase the complexity of the test items. The more complex the items are, the higher is the probability that participants would not have been trained on that particular sentence structure and, therefore, do not have a memorized solution algorithm. On the other hand, in developing the test, we wanted to avoid complex grammar that may be unfamiliar to younger participants and to those who are nonverbal or have intellectual disabilities. We also wanted to avoid tasks involving synthesis of many mental objects that could overwhelm their attention and working memory. Accordingly, we tried to use simple grammatical structures and limit the number of mental objects that had to be arranged into a novel position as much as possible.

With these limitations, there is no perfect single test item to unequivocally assess PFS. At least theoretically, interpretation of any syntactically rigid sentence structure can be routinized into an individual’s implicit memory. If a participant has been trained on a particular item for an extended period of time, any item in the LEPS test can be performed correctly without imagining a novel combination of objects in the process of PFS. Thus, instead of relying on any single item, the LEPS test has 10 items that assess PFS, using various syntactic structures in the hope that most items have not been routinized. Accordingly, the results of the LEPS test have to be interpreted with all the items considered integrally: the combined score of all items is used to assess participant’s PFS ability. The higher the LEPS score, the greater the evidence of developed PFS ability; correct answers in several items on LEPS test shall not be definitively interpreted as an indicator of PFS, especially in individuals with many years of language therapy who could have ingrained interpretation of rigid canonical syntax into context-dependent algorithm. We suggest to use the score of 7 as a minimum evidence for the PFS ability with higher scores providing greater confidence of PFS.

### Why LEPS test includes tangible objects instead of pictures?

For a typical adult, following an instruction to point to a correct picture (e.g., “a whale ate a man” vs. “a man ate a whale”) is no harder than arranging physical toys (a whale and a man) in the correct position. This is not the case for some participants with impairments (Lin & Chiang, 2014; Maljaars et al., 2012). Consider Peter, a 7-year and 7-month-old fully verbal child with ADHD, his case study is described in supplementary material (Case study S1). Peter’s performance on paper-based tests was strikingly different from his performance with physical toys. Peter received a standardized score of 74 on the Fluid Reasoning Index of the WPPSI-IV (i.e. lower than 96% of population; Matrix Reasoning=6, Picture Concepts=5), an IQ test in which the participant has to select a picture that represents the correct answer. This IQ score indicates that Peter has failed all items that examine PFS (Vyshedskiy, Dunn, et al., 2017). We confirmed these observations by testing Peter with our proprietary paper-based test. Peter showed understanding of the concept of matrix analogies by succeeding in all simpler items that required “finding the same objects” and “integration of color, size and number modifiers.” But upon being asked to point to a picture depicting “the man ate the whale” or “the whale ate the man,” Peter answered randomly. Did Peter understand the difference between “the man ate the whale” versus “the whale ate the man?” Although Peter’s performance on the paper-based test was below the chance level, his performance increased to 100% accuracy when he was allowed to show his answer with physical objects (item 5). In fact, Peter has succeeded in all but one LEPS item (he did not reliably understand the difference between the passive and active forms of the verb “eat”) and received a LEPS total score of 9. The LEPS test was a superior measurement for PFS than the paper-based WPPSI-IV test.

It is tempting to neurologically dissect Peter’s ability to understand the difference ‘who ate who’ with physical objects (item 5) from his failure in the equivalent question in a paper-based test. Clearly, Peter’s Wernicke’s area was capable of comprehending the meaning of words, his Broca’s area was capable of assigning word forms to a grammatical group (such as noun or verb), and his LPFC was capable of purposeful synthesis of disparate objects together (in this case the objects were “the man” and “the whale”). From this, we can speculate that Peter’s ADHD is to blame for his failure in paper-based tasks, since tangible objects have been shown to have a greater influence on attention than objects shown in pictures (Gomez, Skiba, & Snow, 2018). It is likely that use of physical toys captured Peter’s attention on the task much more than paper could.

Peter was re-tested with our proprietary paper-based test 4 months after the initial test. At that time, he was taking 30mg of Ritalin daily. This time Peter answered all paper-based items correctly, including items testing PFS.

Peter’s case is a good demonstration of the dissociation between *attention* and PFS. Both attention and PFS are functions of the LPFC. In neurotypical children, attention and PFS are acquired concurrently. However, the dissociation of attention and PFS may be observed in some participants with impairments. In Peter, PFS was normally developed, while there was severe deficit in his attention (that was later corrected by Ritalin). On the other hand, in most late-first-language-learners, attention was normally developed (Curtiss, 1981) while PFS was not (Vyshedskiy, Mahapatra, et al., 2017).

### LEPS can be used to diagnose PFS paralysis and to monitor PFS acquisition in vulnerable children

The importance of early introduction of language to children is widely recognized (Eldevik et al., 2010; Law & Levickis, 2018; Peters-Scheffer, Didden, Korzilius, & Sturmey, 2011; Virués-Ortega, 2010). Vulnerable individuals include children with congenital deafness, ASD, PDD, and any other children with potential for language delay. Several government laws and regulations aim to identify vulnerable individuals. In 1999, the U.S. Congress passed the “Newborn and Infant Hearing Screening and Intervention Act,” which gives grants to help states create hearing screening programs for newborns. Otoacoustic Emissions Testing is usually done at birth, followed by an Auditory Brainstem Response if the Otoacoustic Emissions test results indicated possible hearing loss. Such screening allows parents to expose deaf children to a formal sign language as early as possible and therefore to avoid any delay in introduction to full *recursive* language. When congenitally deaf children are exposed to full recursive spoken or sign language early, their function of PFS develops normally (Mayberry, 2002).

American Academy of Pediatrics (AAP) recommends universal screening of 18- and 24-month-old children for ASD, and also that individuals diagnosed with ASD begin to receive no less than 25 hours per week of treatment within 60 days of identification (Maglione et al., 2012). Despite the AAP recommendation, two-thirds of US children on the autism spectrum under the age of 8 fail to get even the minimum recommended treatment (Hume, Bellini, & Pratt, 2005) because of major problems with the availability, quality, and general funding for early intervention programs (Bibby, Eikeseth, Martin, Mudford, & Reeves, 2002; Jacobson, 2000; Johnson & Hastings, 2002). Since the AAP’s 2007 recommendation of universal early screening, there has been a sharp increase in demand for ASD-related services (58% on average, Ref. (Wise, Little, Holliman, Wise, & Wang, 2010). However, according to a recent study, most states have reported an enormous shortage of ASD-trained personnel, including behavioral therapists (89%), speech-language pathologists (82%), and occupational therapists (79%) (Wise et al., 2010). In many states children are getting less than 5 hours per week of service (Wise et al., 2010). Families of newly diagnosed children often face lengthy waitlists for therapy, leaving children without treatment during the most critical early period of development. Thirty to forty percent of individuals diagnosed with ASD receive inadequate therapy. This can cause lifelong PFS paralysis resulting in inability to understand spatial prepositions and recursion (Vyshedskiy, Mahapatra, et al., 2017).

Propensity to acquire PFS seems to start before the age of two (Bick et al., 2015), reduces notably after five years of age (Basser, 1962; Boatman et al., 1999; Krashen & Harshman, 1972; Lenneberg, 1967; Pulsifer et al., 2004; Vyshedskiy et al., 2019), and ceases completely after puberty (Vyshedskiy, Mahapatra, et al., 2017). As the result of this strong and short critical period, the ability of children to acquire PFS can be significantly diminished by the time they enter the public school system. Timely identification of PFS acquisition delay could facilitate a more comprehensive understanding of a child’s cognitive strengths and weaknesses, which would in turn lead to a more targeted intervention therapy.

### LEPS test can be used for high-functioning versus low-functioning classroom assignment

PFS may be the most salient function of intellect that distinguishes high-functioning from low-functioning individuals. Indeed, PFS ability enables individuals to understand complex explanations by imagining novel scenarios in their mind. Acquisition of PFS is a watershed moment in a life of a child that leads to understanding of complex explanations, fairy tales, and vast improvements in general learning (Burke & Cerniglia, 1990; Wingate et al., 2014).

Conversely, PFS paralysis results in individual’s inability to understand fairy tales and complex recursive explanation, commonly associated with low-functioning individuals. Consistent with this hypothesis, LEPS test had near perfect prediction power of high-functioning vs. low-functioning classroom assignment, Table 2. All students with LEPS total score ≥7 were assigned by teachers to high-functioning classes. All but two students with LEPS total score <7 were assigned by teachers to low-functioning classes: LEPS class assignment prediction rate was 90%. Consistent with teachers’ sentiment that “IQ score is unreliable for student class assignment,” IQ score was much worse at predicting high-functioning vs. low-functioning classroom assignment. With the score of 65 as a threshold, full-scale IQ class assignment prediction rate was only 50%. PFS and its assessment by LEPS total score were better predictors of child’s educational abilities (as interpreted by teachers) than the IQ score.

### Verbal/nonverbal nomenclature is not informative of PFS ability

Among five nonverbal participants with impairments who were using augmentative and alternative communication devices, one (20%) had demonstrated PFS ability (LEPS score≥7). Among 18 verbal participants with impairments 8 (44%) demonstrated PFS ability (LEPS score≥7). We conclude that verbal/nonverbal definitions are not indicative of individuals’ PFS abilities: both verbal and nonverbal individuals can have high or low PFS.

### Comparison of LEPS to the Token Test for children

The Token Test for children, part five (A. De Renzi & Vignolo, 1962; E. De Renzi & Faglioni, 1978), comes closest to standardized assessment of the PFS ability. However, the Token Test is more complex and, therefore, less sensitive to minimal PFS abilities. Consider, for example the task based on spatial prepositions: the Token Test is using instruction “Put the red circle on the green square;” the LEPS test uses shorter instructions: “Put the lion on the giraffe.” The additional adjectives in the Token Test add several layers of complexity by requiring the subject to 1) parse the sentence into three rules: (1) adjective + noun + (2) spatial preposition + (3) adjective + noun (Broca’s area), 2) store these three rules in working memory (LPFC), 3) integrate the first adjective and the noun (temporal cortex under the executive control of the LPFC), 4) store the result of integration in memory (temporal cortex), 5) integrate the second adjective and the noun (temporal cortex under the executive control of the LPFC), 6) store the result of the second integration in memory (temporal cortex), 7) understand the meaning of the spatial preposition (Broca’s area), 8) combine the two mental objects (temporal cortex under the executive control of the LPFC). Conversely, the shorter instructions of LEPS test (“put the lion on the giraffe”) require fewer neurological steps: 1) parse the sentence: noun + spatial preposition + noun (Broca’s area, only one rule), 2) understand the meaning of the spatial preposition (Broca’s area), 3) combine the two mental objects (temporal cortex under the executive control of the LPFC). Each additional neurological step makes the solution much harder for an individual with unusually-short auditory memory. As a result, interpretation of the shorter LEPS instruction may be significantly simpler to individuals with impairments.

The LEPS test also has simpler vocabulary: 1) LEPS uses fewer spatial prepositions; 2) LEPS uses fewer adjectives; 3) nouns used in LEPS (e.g., animal names) are learned by children at younger age than geometric figures used in the Token Test. Finally, LEPS uses simpler grammar than the Token Test. With greater lexical, grammatical, and working memory demands, the Token Test is more a *receptive language* test, while LEPS uses minimal language to evaluate the PFS ability. E.g., LEPS mental size comparison question “Imagine an *elephant* and a *chicken.* Which one is bigger?” simply invites the subject to recall two familiar animals. There is no grammar or syntactic processing here. This item is based on the observation that in order to compare animal’s relative sizes, they have to be combined together in the same mental frame – the process that relies on PFS. Certainly, mental size comparison questions can be answered without PFS, if a subject has seen a picture of the two animals displayed together or stores the semantic memory of size difference. As a result, this type of questions would not work for adults.

## Limitations

There is no single perfect measurement technique for PFS in children. Simpler tests that rely on common sentence structure (“put the green cup inside the blue cup”) can all be trained into automatic algorithms that do not involve creating any novel mental images. More difficult tests, such as the Tower of London (Shallice, 1982), require significant attention and working memory that is often acquired at an older age. Still other tests, such as 2-digit number multiplication (Zago et al., 2001) are not appropriate since children do not know numbers or the concept of multiplication. The LEPS test attempts to strike a balance between common and complex questions to present children with a task of imagining novel combinations of objects in their mind. Certainly, such approach has its limitations.

First, the LEPS test is not applicable to the most children younger than 2.5 years, as those children are not yet familiar with words for colors, sizes, and spatial prepositions *inside*, *on top of*, and *under.* However, children, even if they were not familiar with some words, can often grasp the meaning of those few words during the test: each object used in LEPS is named and each spatial preposition is explained in a demonstration before the test to avoid testing vocabulary.

Second, performance in the LEPS test depends on a child’s attention and motivation. In this regard the LEPS test is no different from other intelligence tests in which children have to stay focused throughout the test. As discussed above, LEPS’ use of physical objects instead of pictures makes it easier for children with attention deficit disorder and can result in a better measure of their fluid intelligence.

## Conclusions

We describe a 10-item Linguistic Evaluation of Prefrontal Synthesis (LEPS) 5-minute test designed for quick assessment of the most complex component of imagination, Prefrontal Synthesis (PFS). LEPS items use spatial prepositions and non-canonical syntax to present participants with a set of novel questions that participants have never encountered before. The sum of 10 items results in the LEPS total score that ranges from 0 (no PFS ability was demonstrated) to 10 (full PFS ability). Internal consistency of LEPS was good (Cronbach’s alpha = 0.95). LEPS exhibited excellent test–retest reliability, very high inter-observer agreement, good known-group validity, and excellent ability to predict high-functioning vs. low-functioning class assignment in participants with impairments. As LEPS does not rely on productive language, it may be an especially useful tool for assessing the development of minimally-verbal children.

## Acknowledgments

We wish to thank Dr. Max Lianski, and Dr. Leo Fridman for productive discussion. We thank Dr. Petr Ilyinskii for scrupulous editing of this manuscript and Benjamin Vyshedskiy for help with data analysis. We thank Havilah Butsick for her contribution in participants testing.

