## SupplementalMaterial for "Novel Linguistic Evaluation of Prefrontal Synthesis (LEPS) test measures prefrontal synthesis acquisition in neurotypical children and predicts high-functioning versus low-functioning class assignment in individuals with autism"

Developed by Andrey Vyshedskiy, Ph.D, Boston University and [ImagiRation.com](http://imagiration.com/)

Directions: This form is intended to measure LEPS score.

Scoring: For each item, enter the score of 1 if a participant demonstrated an understanding of the instructions by successfully completing at least ¾ of the item’s tasks ($\geq$75% accuracy). Enter a score of 0 if a participant completed less than ¾ of the item’s tasks ($\leq75\%).$ To calculate the total LEPS score, add the scores for each item. Highest possible score is 10.

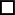
Name of Child:______________ _______________	Female	Age: ________

Last	First	Date of Birth: ________

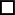
Form completed by: ________________	Male	Today’s Date:________

Language spoken at home: ________________
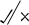

| **1. Integration of Modifiers** |
| --- |
| 1: Give me the *long* *red* straw |
| 2: Give me the *short* *green* straw |
| 3: Give me the *small* *red* Lego |
| 4: Give me the *large* *blue* Lego |
| Score (0/1): |
| **2. Stacking Cups (demonstrate an example and let child repeat after you)** |
| 1: Put the *green* cup inside the *blue* cup |
| 2: Put the *red* cup inside the *green* cup |
| *3:* Put the *green* inside the *orange* cup |
| 4: Put the *orange* inside the *blue* cup |
| Score (0/1): |
| **3. Non-canonical Syntax with Cups** |
| 1: Inside the *blue* cup, put the *green* cup |
| 2: Inside the *red* cup, put the *orange* cup |
| 3: Move the cups so that the *orange* cup is inside the *green* cup |
| 4: Imagine the *green* cup inside the *red* cup. Move the cups to show this |
| Score (0/1): |
| **4. Combination of Plush Animals (demonstrate one animal inside the other and let the child practice)** |
| 1: Show me: the *giraffe* ate the *elephant* |
| 2: Show me: the *lion* ate the *monkey* |
| 3: Show me: the *monkey* ate the *giraffe* |
| 4: Show me: the *elephant* ate the *lion* |
| Score (0/1): |
| **5. Passive Verb Tense with Plush Animals** |
| 1: Show me: the *lion* was eaten by the *giraffe* |
| 2: Show me: the *monkey* was eaten by the *elephant* |
| 3: Show me: the *giraffe* ate the *monkey* |
| 4: Show me: the *elephant* was eaten by the *lion* |
| Score (0/1): |
| **6. Spatial Prepositions with Plush Animals (demonstrate *under* and *on top of* spatial prepositions with animals and let the child practice)** |
| 1: Put the *giraffe* under the *monkey* |
| 2: Place the *elephant* on top of the *giraffe* |
| 3: Put the *lion* on the *elephant* |
| 4: Place the *monkey* under the *lion* |
| Score (0/1): |
| **7. Recursion with Spatial Prepositions and Plush Animals** |
| 1: Put the *monkey* under the *lion* and on top of the *giraffe* |
| 2: Place the *lion* on top of the *giraffe* and under the *elephant* |
| 3: Move the *monkey* so that it is under the *lion* and on top of the *elephant* |
| 4: Put the *elephant* on top of the *giraffe* and under the *monkey* |
| Score (0/1): |
| **8. Mental Size Comparison** |
| 1: Imagine an *elephant* and a *chicken*. Which one is bigger? |
| 2: Imagine a *mouse* and a *cat*. Which one is bigger? |
| 3: Imagine a *lion* and a *cat*. Which one is bigger? |
| 4: Imagine a *chicken* and a *cow*. Which one is bigger? |
| Score (0/1): |
| **9. Mental Reasoning – Animals** |
| 1: If a *monkey* ate a lion, which one is still alive? |
| 2: If a *bird* was eaten by a *cat*, which one is still alive? |
| 3: If a *dog* ate a *cow*, which one is still alive? |
| 4: If a *bear* was eaten by a *mouse*, which one is still alive? |
| Score (0/1): |
| **10. Mental Reasoning – Cups** |
| 1: Imagine the *red* cup inside the *green* cup, which is at the bottom |
| 2: Imagine the *blue* cup inside the *yellow* cup, which cup is on top? |
| 3: Imagine the *blue* cup is inside the *red* cup, which is on the bottom? |
| 4: Imagine the *yellow* cup inside the *green* cup, which is on top |
| Score (0/1): |
| TOTAL LEPS SCORE (0-10): |

**Instructions**

Item 1: Confirm that the participant knows color names of Legos, straws, and cups. For example, point to a red object and ask the participant what color it is, or ask the participant to show you a red item.

Item 2: Ask the participant to identify the color of each of the four cups. For two colors and two colors only, demonstrate what it looks like to put one cup inside of the other. Ask the child to repeat after you. Do not use this color combination in the following tasks.

Item 3: This item does not require a demonstration, however if one is necessary, show the participant how to complete one task.

Item 4: Ask the participant to name the plush animals. Make sure your subject can select all animals after you name them. Demonstrate what it looks like for one animal to “eat” the other by pushing the animal that was eaten into the other animal.

Item 5: This item should not require a demonstration, as it is essentially the same action as item 4. However, if a participant is confused by the instructions, one example can be shown to aid the participant.

Item 6: Demonstrate to the participant what an example would look like, e.g. show them a lion on top of a giraffe and say “this is what it looks like when a lion is on top of a giraffe.”

Item 7: Always use the middle animal as the point of reference, e.g. put the monkey on top of the lion and under the giraffe. Both sets of direction revolve around the monkey.

Items 8 to 10: Instruct the participant that they will not be using any physical objects for these tasks, but that it will all be in their heads. In items 8 to 10, participants must determine the correct answer by using their own mental representations of objects, without the use of physical objects.

**Scoring**

For each task, put either a
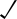
 or an
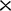
 in the box to the right to note correct or incorrect responses. If necessary, the test administrator can give more than four tasks. Then, for each item, calculate the accuracy (i.e. ¼ = 25%, ^2^/_4_ = 50%, ¾ = 75%, ^4^/_4_ = 100%). Accuracy of 75% and above equals the score of 1, while anything below that equals the score of 0.

Calculate the sum of all items. The highest possible score is 10. A score of 10 indicates that the participant successfully completed each task with at least 75% accuracy and thus received the highest possible score. Similarly, a participant with successful completion of seven items would receive a score of 7, a participant with successful completion of no items would receive a score of 0, and so on.

Case study S1: Peter – a 7 year-7 month-old fully verbal male with ADHD

Peter is a male who exhibits significant difficulty following simple and complex instructions and remaining attentive. He was tested at the age of 7 years and 7 months. He lives in the Boston area with his parents and two brothers. Peter’s mother grew concerned of his clumsiness and short attention span when he was in preschool, and sought evaluations to determine whether it was caused by forgetfulness or an inability to understand instructions. Peter received a diagnosis of Attention Deficit Hyperactivity Disorder (ADHD combined type – both inattentiveness and hyperactivity) at age 6 years 10 months. Peter was also diagnosed with an Unspecified Communication Disorder and a Developmental Coordination Disorder, reflecting his language and motor delays. Despite this, Peter currently plays well with the other kids of his age and has no difficulty making friends. He is extremely talkative, and does not appear atypical or inhibited in the way he interacts with his environment and those around him (e.g. makes eye contact, enjoys being outside and being active, etc.). Peter’s parents describe him as a happy and easy-going boy. However, he fidgets constantly and is easily distracted by outside stimuli. Peter is unable to efficiently complete tasks at school and at home because he becomes easily frustrated when something requires his focus and attention for any extended period of time. Additionally, he exhibits delays in receptive and expressive language development. Most notably, he is unable to understand complex directions and questions, and responds to questions with short, simple sentences.

The Wechsler Preschool and Primary Scale of Intelligence, Fourth Edition (WPPSI-IV) was administered to measure Peter’s cognitive abilities. He received a standardized IQ score of 79 (8^th^ percentile), which falls within the Borderline range. This indicated a clinically significant deficit in overall intellectual abilities. Within the WPPSI-IV, Peter received a score of 74 on the Fluid Reasoning Index (4^th^ percentile; Matrix Reasoning=6, Picture Concepts=5), which measured nonverbal reasoning skills. The Wechsler Individual Achievement Test, Third Edition (WIAT-III) was used to measure Peter’s educational performance. The test results indicated that most of Peter’s educational abilities are at the Kindergarten level, slightly below a normal level for his age. The results of this IQ measures, as well as clinical observations of Peter’s behavior, was in line with his mother’s reports that he struggles with behavioral inhibition, attention, and certain aspects of executive function. On the parent-reported Vineland-III, Peter received a standardized Communication score of 75 (5^th^ percentile)). Peter does not yet meet criteria for an intellectual disability; however, his ADHD severely inhibits his performance at school. At the time of LEPS test, ADHD was diagnosed, but Peter was not taking any medications yet.
